# THE DYNAMICS OF HISTORICAL AND RECENT RANGE SHIFTS IN THE RUFFED GROUSE (*Bonasa umbellus*)

**DOI:** 10.1101/2020.08.23.263194

**Authors:** Utku Perktaş

## Abstract

Climate variability is the most important force affecting distributional range dynamics of common and widespread species with important impacts on biogeographic patterns. This study integrates phylogeography with distributional analyses to understand the demographic history and range dynamics of a widespread bird species, the Ruffed Grouse (*Bonasa umbellus*), under several climate change scenarios. For this, I used an ecological niche modelling approach, together with Bayesian based phylogeographic analysis and landscape genetics, to develop robust inferences regarding this species’ demographic history and range dynamics. The model’s predictions were mostly congruent with the present distribution of the Ruffed Grouse. However, under the Last Glacial Maximum bioclimatic conditions, the model predicted a substantially narrower distribution than the present. The predictions for the Last Glacial Maximum also showed three allopatric refugia in south-eastern and west-coast North America, and a cryptic refugium in Alaska. The prediction for the Last Interglacial showed two separate distributions to the west and east of the Rocky Mountains. In addition, the predictions for 2050 and 2070 indicated that the Ruffed Grouse will most likely show slight range shifts to the north and will become more widely distributed than in the past or present. At present, effective population connectivity throughout North America was weakly positively correlated with F_*st*_ values. That is, the species’ distribution range showed a weak isolation-by-resistance pattern. The extended Bayesian Skyline Plot analysis, which provided good resolution of the effective population size changes over the Ruffed Grouse’s history, was mostly congruent with ecological niche modelling predictions for this species. This study offers the first investigation of the late-Quaternary history of the Ruffed Grouse based on ecological niche modelling and Bayesian based demographic analysis. The species’ present genetic structure is significantly affected by past climate changes, particularly during the last 130 kybp. That is, this study offers valuable evidence of the ‘expansion–contraction’ model of North America’s Pleistocene biogeography.

## INTRODUCTION

The Pleistocene’s glacial periods dramatically affected the biological diversity of the temperate region of the Northern Hemisphere. For instance, widespread temperate species had to find climatically favorable places to survive during the harshest periods of last glaciation episode, which peaked in approximately 22-26 kybp (Clarck et al. 2009). Evolutionary biologists, who have long been interested in the relationship between earth history and biogeographic processes (e.g. vicariance and dispersal), have explained this in terms of the glacial refugia hypothesis.

Numerous phylogeographic studies of North American bird species have tested the hypothesis to explain the geographical structure, demographic history, and gene flow of widespread bird species (Mila et al. 2000, Zink et al. 2001, Mila et al. 2006, Mila et al. 2007, Barrowclough et al. 2011, Pulgarin-R and Burg 2012, van Els et al. 2012), as well as subspecies distribution and speciation (Klicka et al. 2011, Barrowclough et al. 2018). In general, these studies show that many common North American bird species used locations in the south of the continent as a refugium during the Last Glacial Maximum. However, several recent phylogeographic studies that included past distribution projections of various bird species have focused on cryptic refugia located in northern North America (e.g. van Els et al. 2012).

The present study focused on mitochondrial sequences and occurrence records of the Ruffed Grouse (*Bonasa umbellus*), a common and widespread North American bird species. These data were used to examine whether the Pleistocene glaciations affected the demography of Ruffed Grouse populations.

Several features make the Ruffed Grouse an appropriate model organism for this purpose. First, this species is associated with variety of climax forest types, including temperate coniferous rain forest and relatively arid deciduous forest, which were mostly covered by ice during the Last Glacial Maximum. Hence, its current widespread distribution throughout North America (Fig. 1) suggests that the origin of populations from northern North America must be one or more glacial refugia in the south of its distribution range. Second, the Ruffed Grouse is not considered migratory bird species though it exhibits some seasonal variations in mobility (Johnsgard 1983). Various studies report that its dispersal capacity is generally limited, without large differences between different age groups within the species (Chambers and Sharp 1958, Hale and Dorney 1963). Third, since the species is highly dependent on its environment, it is reasonable to expect a balance between climate and its current distribution; and, if so, this assumption should also be historically valid (i.e. for the Last Glacial Maximum). It is thus crucial to test how the Ruffed Grouse responds to past and future climate change scenarios to understand its demographic history and the dynamics of its range shifts. Finally, the Ruffed Grouse is also an appropriate model organism because its mitochondrial phylogeographic structure is well studied. Two recent mitochondrial DNA (mtDNA) studies have analyzed intra-species gene variation for two different mitochondrial genes (Jensen et al. 2019, Honeycutt et al. 2019). Jensen et al. (2019) focused on landscape effects on the genetic structure of Ruffed Grouse; Honeycutt et al. (2019) focused on mtDNA variation using cytochrome-*b* (cyt-*b*) gene across almost the complete distribution range.

**Figure 1.**
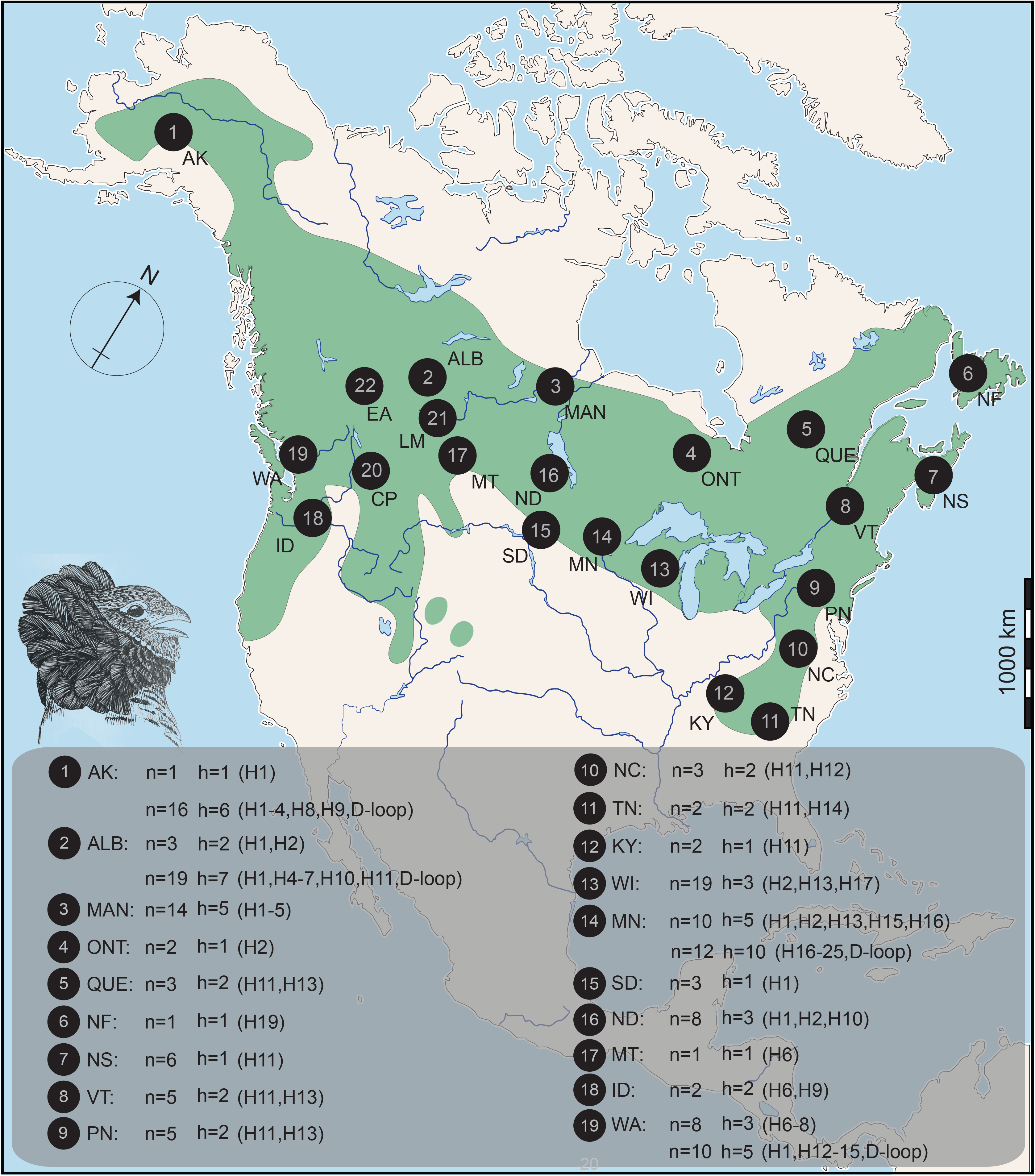
Ruffed Grouse distribution based on Johnsgard (1983) and Madge & McGowan (2002). Map showing locations, sampling size(n), and haplotype numbers (h).

Distributional analyses (e.g. ecological niche modelling) is an important methodological development designed to predict past, present and future geographic distribution of the species. Its integration with phylogeography often incorporates geographic diversity patterns into genetic diversity and diversity analysis, providing strong inferences for the demographic history of species (Carstens and Richards 2007, Alvarado Serrano and Knowles 2014). Because neither study discussed the species’ demographic history in detail, the present study integrates phylogeography with distributional analyses to understand the demographic history and range dynamics of Ruffed Grouse under climate change scenarios. Hence, this study extends and integrates the work of Jensen et al. (2019) and Honeycutt et al. (2019) through novel analyses of demographic history and distributional projections derived from ecological niche models (ENMs).

## MATERIAL and METHODS

### Ecological Niche Modelling

Input data – I analyzed species occurrence data from e-Bird (www.ebird.org), ranging from 2000 to 2019 (n = 267027), before checking for sampling bias and spatial autocorrelation (Brown 2014) for occurrence records. I spatially filtered all records to eliminate multiple records, leaving single 25-km records across the species’ distribution. For this, I used the dispersal capacity of the Ruffed Grouse (see Johnsgard 1984 for details). This yielded 4,566 unique occurrence records for ecological niche modelling.

I downloaded bioclimatic data from the WorldClim database (Hijmans et al. 2005, http://www.worldclim.org) for three global climate models (CCSM4, MIROC-ESM, and MPI-ESM-P) for the Last Glacial Maximum (~22 kybp), the mid-Holocene (~6 kybp), the present (~1960-1990), and future conditions based on the rcp45 greenhouse gas scenario (2050 and 2070) at a spatial resolution of 2.5 arc-minutes. Bioclimatic data included 19 bioclimatic variables derived from monthly temperature and precipitation values. Since the Ruffed Grouse is a widespread and common bird species in North America (Fig. 1), all variables were masked to include all North America (−170° to 13° W and −50° to 84° N). I then inspected correlations between these bioclimatic variables to produce three different climatic data sets based on different inter-variable correlation coefficients (0.7, 0.8, and 0.9). These included annual mean temperature and precipitation (BIO1 and BIO12), mean diurnal range (BIO2), isothermality (BIO3), temperature and precipitation seasonality (BIO4 and BIO15), annual temperature range (BIO7), warmest quarter precipitation (BIO18), wettest quarter mean temperature (BIO8), driest quarter mean temperature (BIO9), and driest month precipitation (BIO14).

Modeling – I used the kuenm package (https://github.com/marlonecobos/kuenm; Cobos et al., 2019) in R 3.6.1 (R Core Team, 2019) for all modelling. For this, I used the maximum entropy machine learning algorithm in MaxEnt version 3.4.0 (Phillips et al. 2006, Elith et al. 2011) to model Ruffed Grouse ecological niches for past (Last Glacial Maximum and Holocene), present, and future bioclimatic conditions because MaxEnt mostly performs better than comparable methods (Elith et al. 2006, Wisz et al. 2008).

I ran MaxEnt with the following four steps: (1) A minimum convex polygon (M area) was created from the occurrence records, applying 100 km buffer zones based on the species assumed natural history (Johnsgard 1984). (2) I used nine different regularization multipliers (0.1, 0.2, 0.5, 0.8, 1, 2, 5, 8, 10) and five different feature types, linear (L), quadratic (Q), product (P), threshold (T), and hinge (H), with 29 combinations of the feature types for model calibration. This produced different candidate models for each regularization multiplier and feature type combination. (3) I selected the best candidate model using the Akaike Information Criterion, corrected for small sample sizes (Hurvich and Tsai 1989). I then used partial ROC to conduct the significance tests (Peterson et al. 2008) and evaluated model performance using a 5% training presence threshold to evaluate omissions (Peterson et al. 2011) for four different climatic data sets. (4) I selected the best calibration based on three different statistics before running MaxEnt to produce the models with ten replicates and bootstrap run type. Model outputs were converted to binary predictions based on the 10-percentile training presence thresholding approach (Perktaş et al. 2017) to project the final (‘best’) models (Freeman et al. 2019).

### Landscape Genetics

To characterize population connectivity in the present I inverted the prediction under present bioclimatic conditions for use as a friction layer; i.e. areas of high suitability were converted to areas of low dispersal cost (low resistance areas). I then calculated least-cost corridors (LCCs) among geographic localities sharing haplotypes in three (high, mid, low) classes, using the ‘percentage of least-cost path (LCP) value’ method. LCC class percentages of 5-2%, 2-1% and < 1% were selected, and LCC class weights of 1, 2, and 5 were applied to high, mid, and low classes, respectively. All weighted LCCs were summed to create the dispersal network (Chan et al. 2011, Brown 2014). Except where otherwise indicated, I conducted all analyses using SDMtoolbox version 2.2 (Brown et al. 2017), implemented in ArcGIS version 10.2.2.

I also tested isolation-by-resistance for the complete Ruffed Grouse distribution range to understand how landscape affected genetic differentiation between populations. For this, I used a paired Mantel test, and calculated the correlations between the matrices of genetic distance (*F*_ST_) based on cyt-*b* gene versus resistance values, and between the matrices of LCP costs versus LCP distances. The paired Mantel test was performed with 10,000 random permutations implemented in XLStat version 2019.3.2 (Addisoft). The genetic distance matrix between 19 Ruffed Grouse populations was estimated with DNAsp version 6.12.01 (Rozas et al. 2017). A resistance matrix between 19 populations was created from the LCC map by weighting the distance of each LCC by the resistance values along the corridor.

### Demographic History

I applied the extended Bayesian Skyline Plot (EBSP) method, implemented in BEAST 2 (Bouckaert et al. 2014), to explore the Ruffed Grouse demographic history. This analysis uses coalescent approaches to estimate effective population size change through time. Since earlier studies have not reported any structure (Jensen et al. 2018, Honeycutt et al. 2019), I combined all mtDNA haplotypes for each locus before running the EBSP analysis. This approach made the historical demography results more comparable with the ENM results (e.g. Perktaş et al. 2019). Before the EBSP runs, the best-fit substitution models were identified for both mtDNA loci in MEGA X (Kumar et al. 2018). These were the Tamura and Nei (1993) with a discrete Gamma distribution (TN93 + G, AICc = 1832.275) for the D-Loop, Hasegawa, Kishino and Yano (HKY, AICc = 2123.177) for the cyt-*b* gene. I used both mtDNA loci (D-Loop and cyt-*b*) in one EBSP analysis. Multiple independent extended Bayesian skyline plot runs were performed using the following parameters: linear models, 100 million steps, parameters sampled every 10,000 steps, and a burn in of 10%. For the D-Loop, I used the strict clock model with a default mutation rate under normal prior distribution, and allowed the analysis to estimate the rates relative to the cyt-*b* gene [for birds, the widely-used 2% substitutions /site /million years (Brito 2005, Pereira and Baker 2006), and for grouse species, 5.04% substitutions /site/ million years (Arbogast and Slowinski 1998)]. I then calculated the expansion time using the two different mtDNA mutation rates, i.e. 2% and 5.04% substitutions /site/ million years, as the mutation rate of the mtDNA cyt-b gene. The effective sample size values of the parameters were over 200 for each run.

## RESULTS

### Ecological Niche Modelling and Landscape Genetics

I evaluated 783 candidate models using combinations of 29 feature classes, 9 regularization multipliers, and 3 climatic data sets. The best model for the Ruffed Grouse was provided by the first climatic data set (Set1: BIO1, BIO7, BIO8, BIO12, BIO15), which was significantly different from random (P < 0.001), and met the ≤5% omission criteria set. The model had a regularization multiplier of 0.2 and included linear and product feature classes. Projections for past, present, and future performed better than a random prediction (training AUC = 0.663, sd = 0.0015). The small SDs for the mean AUCs suggested that the model performance was robust to variations in the selection of training occurrence records. Two bioclimatic variables contributed the most to the model (together 65%): annual mean temperature (BIO1, 27.1%) and temperature annual range (BIO7, 37.9%). That is, the Ruffed Grouse uses an environmental space characterized by annual mean temperature (−5°C-15°C) and annual temperature range (Supplement 1).

Under present bioclimatic conditions, the model’s predictions were mostly congruent with the Ruffed Grouse’s present and recent historical distribution (see Fig. 2; for the present distribution, see BirdLife International, 2016). The model primarily predicted areas of high suitability across suitable habitats for the species in North America. Under mid-Holocene bioclimatic conditions, the prediction showed little difference from the actual present distribution (Fig. 2). However, under the Last Glacial Maximum bioclimatic conditions, the model predicted a substantially narrower distribution than the present and mid-Holocene (Fig. 2). Interestingly, predictions for the Last Glacial Maximum included three allopatric refugia in south-eastern and west-coast North America, and a cryptic refugium in Alaska (see Supplement 2). For 2050 and 2070, the model predicted that the range will most likely shift slightly northward with a wider distribution than either the past or present (Fig. 2).

**Figure 2.**
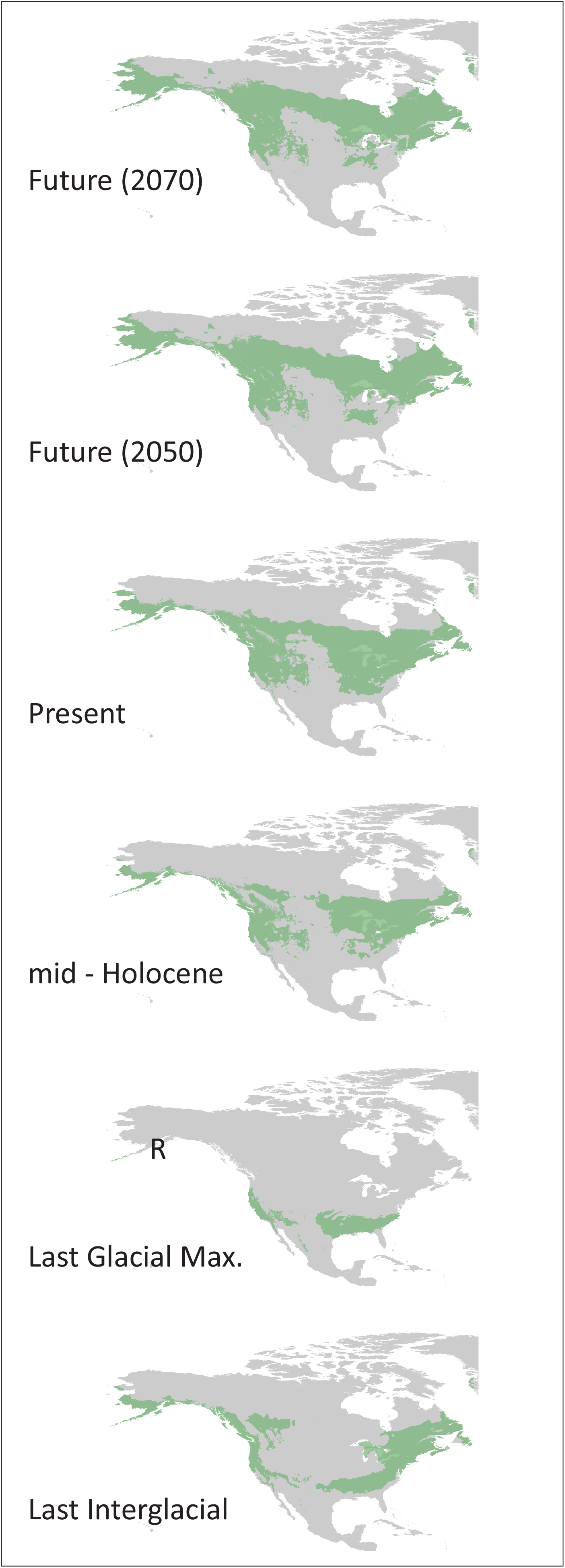
Summary of projections [Last Interglacial, Last Glacial Maximum, mid-Holocene, present, and future (based on 2050 and 2070)] of an ecological niche model for the Ruffed Grouse. The predictions show the 10 percentile training presence logistic threshold results. Predicted species range is shown in green. Individual models of Last Glacial Maximum showed a cryptic refugium in Alaska (R), and they can be seen in Supplement 2.

At present, there is an effective population connectivity throughout North America between populations located in central North America (Fig. 3). Between Alaska and central North American localities specifically, no suitable dispersal corridors appeared to indicate possible isolation-by-resistance. However, the relationship between genetic distance and resistance did not show a strong positive correlation (r = 0.189, P = 0.009). I therefore conclude that there is weak isolation-by-resistance over the species' distribution range. In addition, the relationships between LCP distance and LCP cost showed a strong positive correlations (r = 0.976, P < 0.0001, Fig. 4).

**Figure 3.**
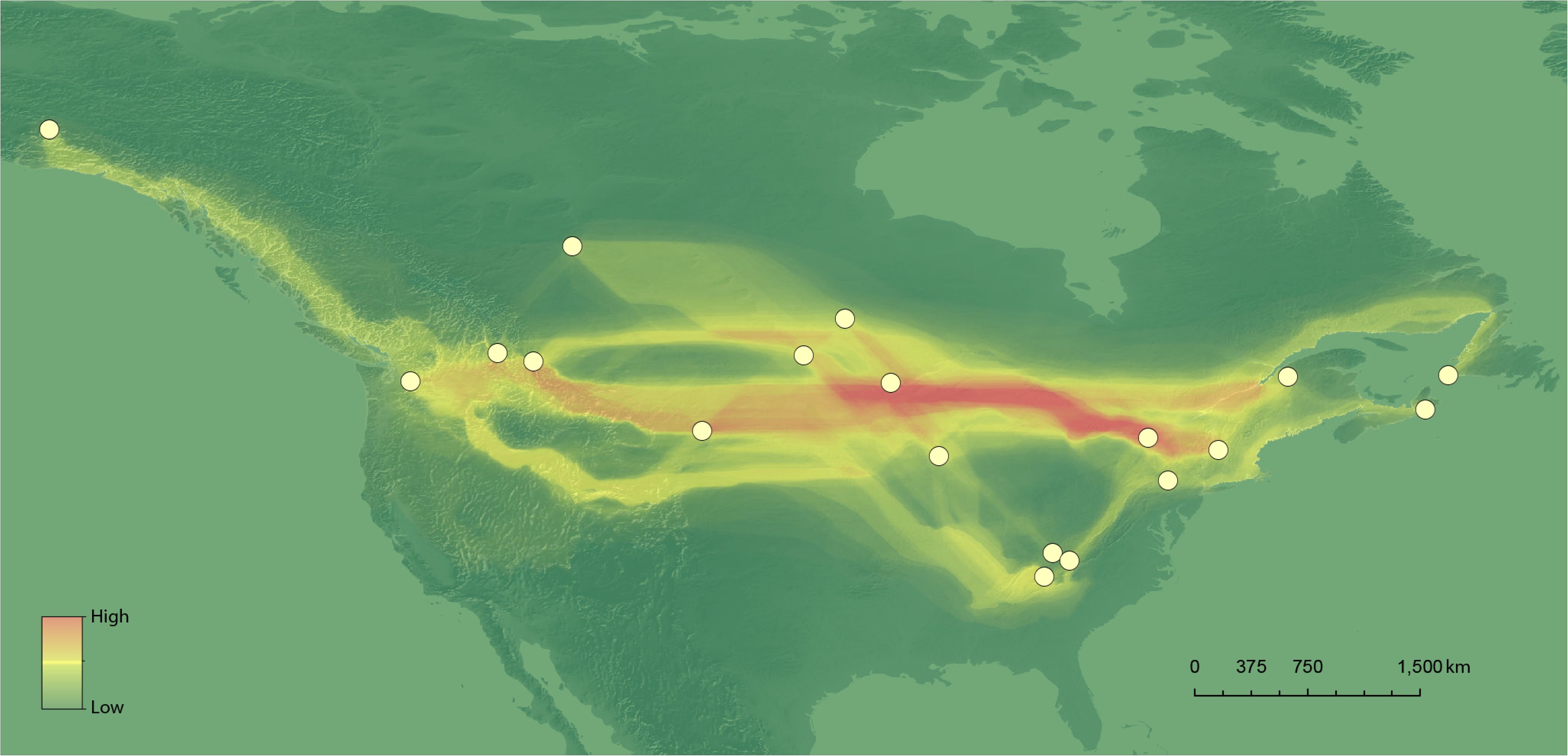
Effective population connectivity between Ruffed Grouse populations. Warmer colors indicate lower inter-population resistance.

**Figure 4.**
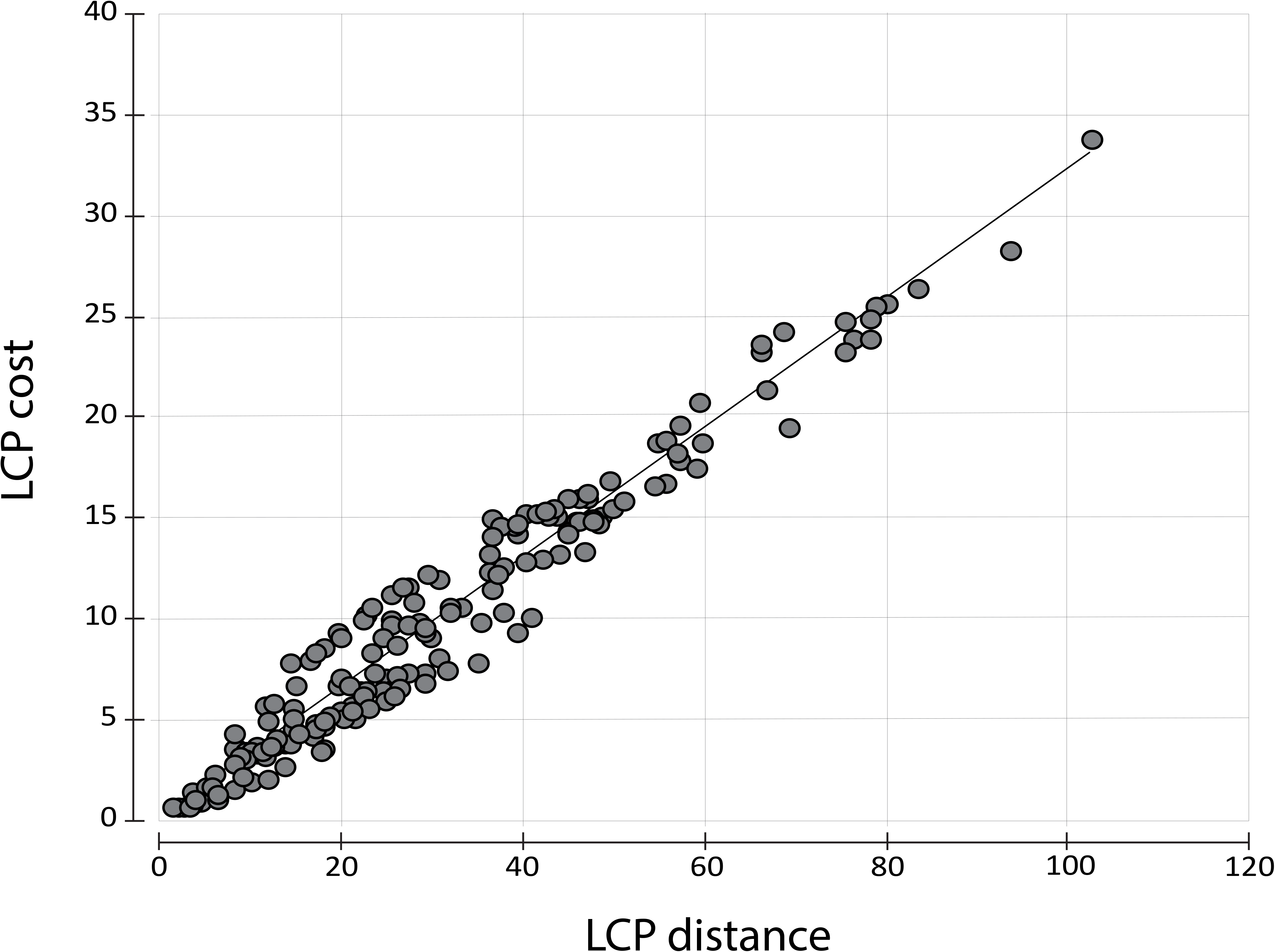
Correlations between **least-cost path (LCP)** distance and **least-cost path** cost.

### Demographic History

Based on a strict molecular clock, D-Loop had a significantly higher substitution rate (mean 7.59% substitutions/ site/ million years when 2% was used for cyt-*b*; mean 18.36% substitutions/site/ million years when 5.04% was used for cyt-*b*) than cyt-*b*. For both rates, the EBSP results provided good resolution of the effective population size changes over the Ruffed Grouse history (Fig. 5). The EBSP indicated a recent sharp demographic expansion since the Last Glacial Maximum (approximately after 20 kybp).

**Figure 5.**
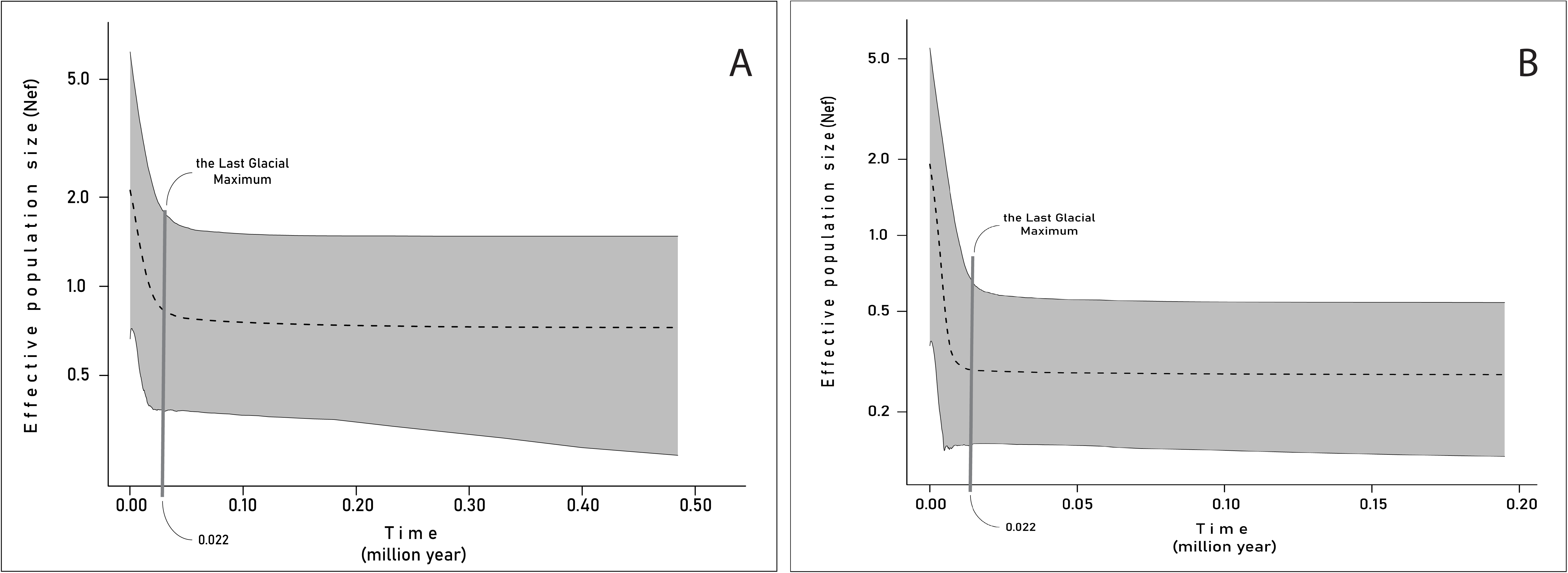
Median effective population size changes over the Ruffed Grouse’s species history. A and B show the results based on 2% and 5.04% divergence rate, respectively. Thin lines indicate 95% highest posterior density interval.

## DISCUSSION

Integrating ecological niche modelling, the associations between environmental variables and a species’ known occurrences can be used to define distribution predictions based on the abiotic conditions within which populations can be maintained (Guissan and Thuiller 2005). Phylogeography usually provides a unique original perspective for robust evaluations of inferences of a species’ demographic history (Knowles, Carstens and Keat 2007, Perktaş and Gür 2015). Several studies on birds that specifically coupled mtDNA phylogeography with ecological niche modeling (Dai et al. 2011, Zhao et al. 2012, Wang et al. 2013, Hung et al. 2013, Zink 2015, Barrowclough et al. 2019) have provided novel insights to understand the impact of climate-driven range shifts in the late Quaternary (i.e from 130 kybp to the present).

In this study, I therefore integrated published mtDNA phylogeography with ecological niche modelling to evaluate the demographic history, including past distributional projections, for a widespread North American bird species, the Ruffed Grouse that is mostly attached to a wide variety of climax forest community type. I also estimated the species’ future range shifts to infer the possible effects of the Anthropocene (Monsarrat, Jarvie and Svenning 2019).

The past distributional predictions indicated allopatric ranges for both the Last Interglacial and the Last Glacial Maximum. The predicted Last Glacial Maximum range was one of the described biogeographical patterns in this common and widespread bird species in North America. It indicated different refugia to the east and west of the Rockies (e.g. American Redstart, *Setophaga ruticilla*, Colbeck et al. 2008). Another example was the refugium in Alaska (e.g. Sharp-tailed Grouse, *Tympanuchus phasianellus*, Perktaş and Elverici, 2019; Canada Jay, *Perisoreus canadensis*, van Els et al. 2012). This type of biogeographic pattern was also valid for a wide variety of climax forest type. Each species (e.g. the Chesnut, the Maple, the White Pine, the Hemlock) in these forest types had also a unique migratory history after the Last Glacial Maximum, including different refugia, and different migration speeds (Pielou 1991).

The current results show that all three refugia from the Last Glacial Maximum were involved in forming the present distribution of the Ruffed Grouse. In addition, the other result suggests that the Last Interglacial played a substantial role in the early differentiation in Ruffed Grouse mtDNA sequences because a clear break between east and west dated back to 130 kybp.

According to ecological niche modelling results, the main breaks at that time were obviously the Rocky Mountains and the Ozark Plateau (Zeisset and Beebee 2008). During the Last Interglacial, the Pacific coast population was evidently restricted to the coast from California to Alaska while the Olympics, Coastal, Cascades, and Sierra Nevada were not effective barriers. The west-coast and Alaskan populations were separated from the east by the Rockies. During the Last Glacial Maximum, the northwestern Rockies, Olympics, Coastal and Cascades served as a barrier separating coastal populations in the west from the north to the south, and forming two refugia in the species’ western range.

The present genetic differentiation between Alaska and the west coast of North America possibly started in the Last Glacial Maximum. This differentiation has been maintained through isolation (i.e. isolation-by-resistance) in the present, as suggested by Jensen et al. (2019). However, the present relationship between genetic distance and resistance only showed a weak positive correlation, which indicates either almost no limit to gene flow across most of the distribution range, especially in the central and eastern regions. Based on mitochondrial D-Loop and cyt-b genes, the Ruffed Grouse is not phylogeographically structured. Based on the cyt-b gene, Honeycutt et al. (2019) found four different haplogroups, whose distributions matched with refugia distributions in the Last Glacial Maximum. Jensen et al. (2019) reported similar results for the western part of the distribution range. At least three high-frequency haplotypes had a geographically structured distribution that matched with the western refugia (i.e. Alaska and the west coast). Both studies made similar phylogeographic inferences that almost all haplotypes were closely related, although some common haplotypes were geographically structured (Avise’s phylogeographic category III; see Avise 2000). This phylogeographic result indicates low or moderate historical gene flow between Ruffed Grouse populations that were not tightly connected historically (e.g. in the Last Glacial Maximum and Last Interglacial; see Perktaş et al. 2019). Due to isolation-by-resistance (Jensen et al. 2019), this conclusion confirms that contemporary gene flow in the western part of the species’ range has been low enough to promote genetic divergence between west-coast and Alaskan Ruffed Grouse populations.

Since both genes showed similar phylogeographic patterns, I used two different mtDNA data sets together to calculate the effective population size changes over the species’ history. The extended Bayesian Skyline Plot analysis showed a substantial population increase after the Last Glacial Maximum The species has reached its present distribution gradually since the late Holocene. Thus, based on my findings for both genes, the species’ demographic history supports the ecological niche modelling results. In contrast to research on other grouse species (e.g. Sharp-tailed Grouse), I found no evidence of a large refugium in southern North America. However, like Sharp-tailed Grouse, this species almost has completely changed its distribution since the last glacial period (Perktaş and Elverici 2019).

Several populations located in Alberta, Manitoba, Ontario, Minnesota, and North and South Dakota contain haplotypes of different clades, indicating potential contact zones and mitochondrial introgression. This suggests that all three refugia influenced the formation of the Ruffed Grouse’s present genetic structure.

This study provides the first investigation of the Ruffed Grouse’s late-Quaternary history based on ecological niche modelling and Bayesian-based demographic analysis. I found that the species’ present genetic structure has been significantly affected by past climate changes, particularly during the last 130 kybp. This study thus provides valuable evidence for the ‘expansion–contraction’ model of North America’s Pleistocene biogeography. In particular, it indicates that it may be more complex than previously thought.

## Supporting information

View of Ruffed Grouse distribution in geographic and environmental space. The top figure shows occurrence records on the map. The bottom figure plots

Three projections of Last Glacial Maximum (A-CCSM4, B-MIROC-ESM and C-MPI-ESM-P).

## ACKNOWLEDGEMENTS

Can Elverici kindly prepared the base map for the ecological niche modelling analyses. Liviu Parau improved the manuscript. Logistic support for the ecological niche modelling analysis was provided by a research project supported by Hacettepe University (project number: FHD-2018-17059). For GenBank accession numbers of sequences that I used in this study, see Jensen et al. 2019 (MK603980–MK604036), and the additional file 2 in Honeycutt et al. 2019.

---

**Supplement 1.** View of Ruffed Grouse distribution in geographic and environmental space. The top figure shows occurrence records on the map. The bottom figure plots known occurrences in a space summarizing annual mean temperature and temperature annual range. Dots with three different colors show G (geographic space), M (dispersal potential of the species), and E (actual species occurrence records).

**Supplement 2.** Three projections of Last Glacial Maximum (A-CCSM4, B-MIROC-ESM and C-MPI-ESM-P).

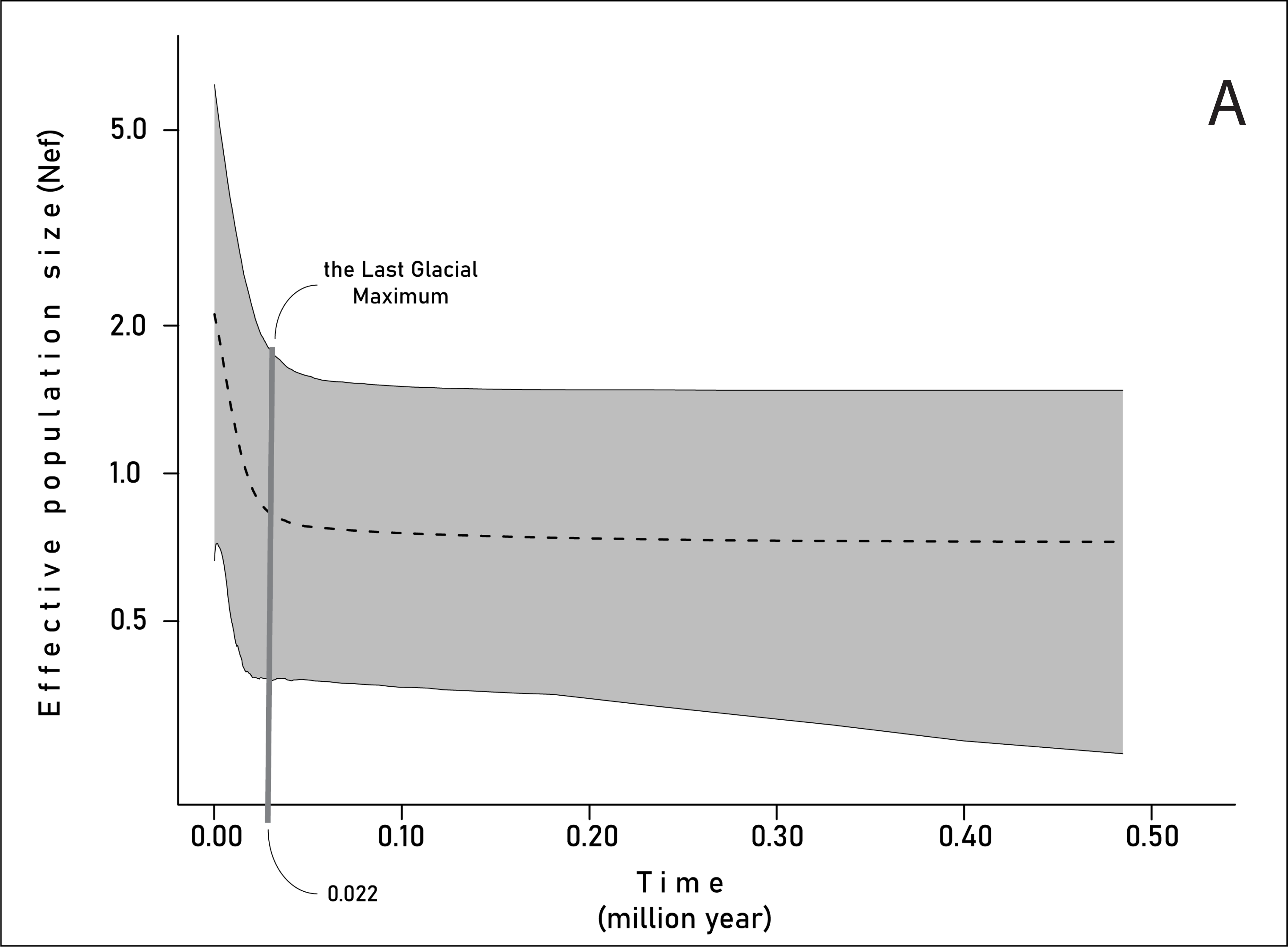

