## Supplementary figures and images for "THE DYNAMICS OF HISTORICAL AND RECENT RANGE SHIFTS IN THE RUFFED GROUSE (*Bonasa umbellus*)"

### Three projections of Last Glacial Maximum (A-CCSM4, B-MIROC-ESM and C-MPI-ESM-P).

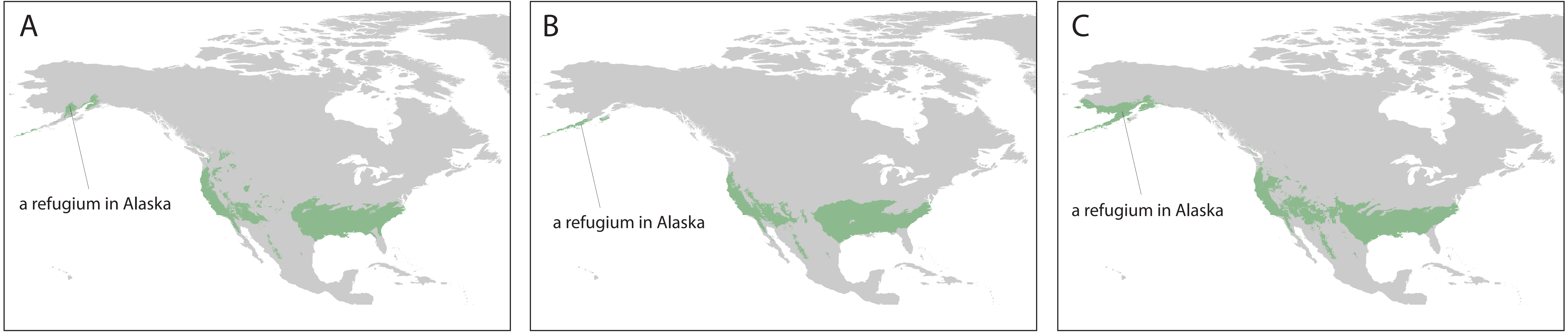

### View of Ruffed Grouse distribution in geographic and environmental space. The top figure shows occurrence records on the map. The bottom figure plots

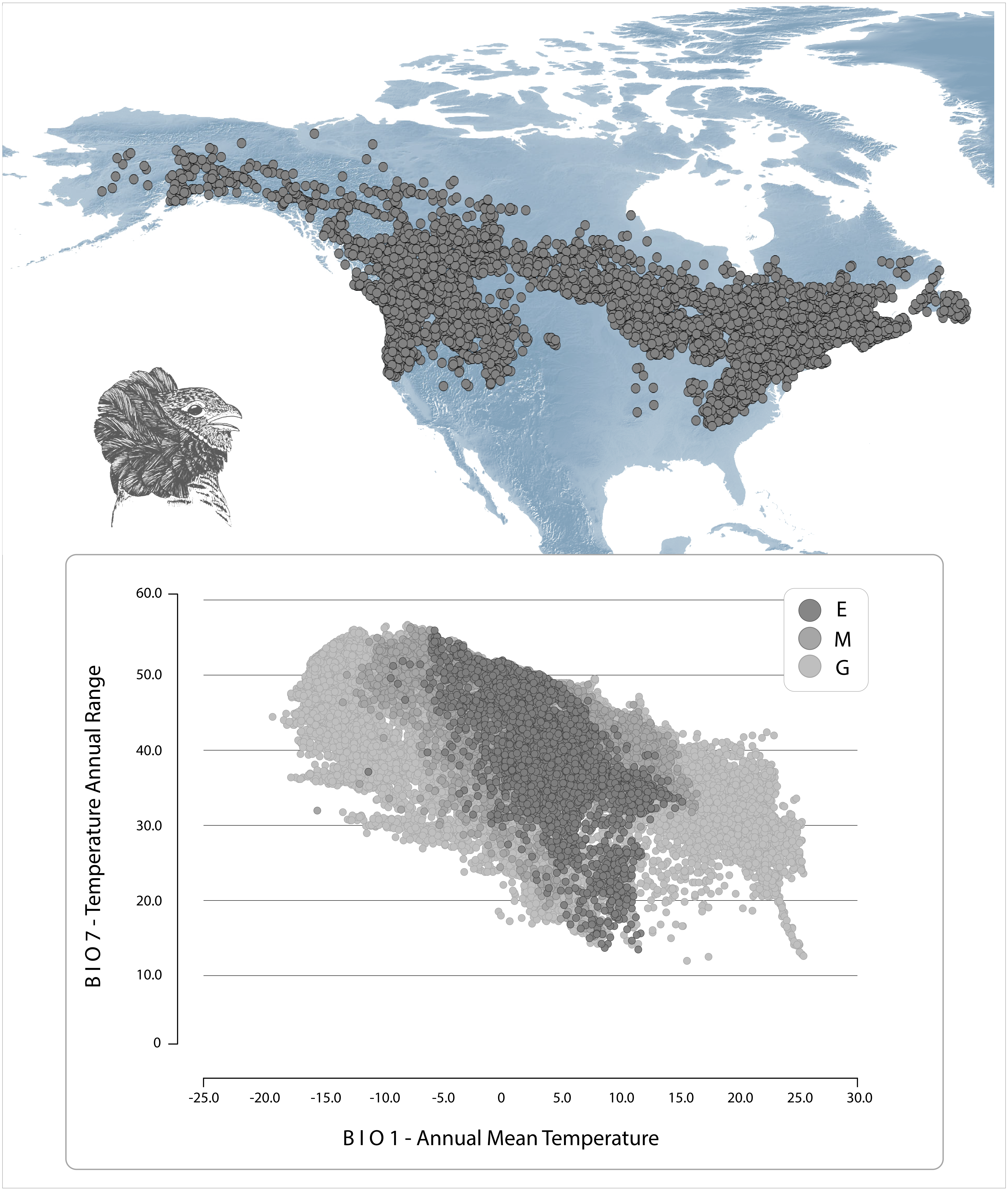
